# Changes in peripheral sensory afference do not alter predictive motor planning: evidence from carpal tunnel syndrome

**DOI:** 10.64898/2026.04.01.715947

**Authors:** Thomas Jacob, B K Mohamed Ibrahim, G Vishnu Babu, K Shyamnath Krishna Pandian, G Karthikeyan, R Krishnamoorthy, K Sridhar, Jahir Hussain, C S Ezhilavan, Sridhar Rajagopal, Sivakumar Balasubramanian, SKM Varadhan

**Author notes:** Corresponding Author: Varadhan SKM, Department of Applied Mechanics and Biomedical Engineering, Indian Institute of Technology Madras, Chennai - 600036 Tamil Nadu.

## Abstract

Anticipatory control organises motor output prior to predictable perturbations and is expressed in multi-digit tasks as anticipatory synergy adjustments (ASAs), which coordinate digit forces before movement onset. Whether such feedforward coordination depends on peripheral sensory input remains unclear. Carpal tunnel syndrome provides a model of altered median nerve afference with within-subject restoration following surgical decompression.

We quantified ASA onset and amplitude in eleven individuals with carpal tunnel syndrome performing a multi-finger grasp-and-release task before and three weeks after decompression surgery. Postoperatively, sensory function improved, and total grip force decreased significantly across task phases, indicating more efficient force regulation. In contrast, ASA onset timing and amplitude were unchanged. Equivalence testing confirmed that pre- and post-operative ASA measures fell within predefined bounds of practical equivalence.

These findings demonstrate a central-peripheral dissociation: feedback-mediated grip force scaling is sensory-dependent and rapidly recalibrates following afferent restoration, whereas feedforward synergy coordination remains stable despite months of degraded peripheral input. The preserved ASA suggests that central motor planning circuits maintain anticipatory coordination through efferent copy or cerebellar-mediated internal models that do not require continuous peripheral recalibration. This resilience may reflect the brain’s ability to maintain predictive motor planning despite chronic sensory degradation, with implications for understanding compensatory mechanisms in peripheral neuropathies and the limits of sensory-dependent motor adaptation.

## 1. Introduction

Voluntary actions typically exhibit premovement cortical activity (Shibasaki & Hallett, 2006). Even the most basic reflexes are subject to modulation by the brain, as descending supraspinal pathways from higher centres can alter their magnitude and timing (Peirs et al., 2020; Sherrington, 1906). Anticipatory control represents a fundamental mechanism in motor control theories, where the nervous system predicts and prepares for future movement demands before they occur (Kao et al., 2021). This feedforward control strategy allows the motor system to make preparatory adjustments that minimise the adverse effects of predictable perturbations or movement requirements (Maffei et al., 2017).

Movements are organised through coordinated coactivation of downstream neurons and actuators by hierarchically higher connected neurons (Santello et al., 2013). This coordinated movement, which includes time-sensitive coactivation and error correction between different individual actuators, is accomplished through synergies. Synergies represent multi-element systems where the nervous system coordinates groups of effectors whose co-variation stabilises some task-relevant performance variables (Santello & Soechting, 2000). The synergistic control brings a fresh perspective to the classical problem of motor redundancy, which puzzled movement scientists about how the nervous system controlled such a complicated physiological system with so many parameters to control. It replaces the redundancy with a principle of abundance, such that multiple equivalent solutions are intentionally preserved and organised via synergies to keep task outcomes stable yet adaptable (Latash, 2012). Synergies enable learned movements to be executed in a seemingly automatic and systematic manner.

Before a planned, quick, or potentially destabilising movement, the central nervous system changes the organisation of synergies, which is known as anticipatory synergy adjustments (ASAs). Their primary goal is to prepare the body for quick action by momentarily decreasing the stability of specific performance variables that are regulated by these synergies. In healthy participants, destabilisation in synergy occurs in anticipation of a new state or position, typically 100-300ms before movement onset (Kim et al., 2006). This anticipatory mechanism reflects feedforward control processes where covariation among elemental variables (such as finger forces) changes in preparation for producing a change in performance variables without immediately altering the magnitude of those variables (Olafsdottir et al., 2005). These are quantified as a reduction in this synergy index before movement onset, indicating a transient reduction in the stability of the performance variable to allow rapid change (Goodman & Latash, 2006). The ASA phenomenon has been consistently observed across various motor tasks, including multi-finger force production, prehensile actions, and postural control. This declines with age (Olafsdottir et al., 2005, 2007) and disease (Falaki et al., 2016; Park et al., 2012, 2014; Rearick et al., 2002).

Carpal Tunnel syndrome is a neuropathy affecting the median nerve due to its compression at a physiologically narrow space in the wrist known as the Carpal Tunnel. This results in slower nerve conduction and consequent sensory deficits, thereby affecting hand function (Padua et al., 2016). Carpal tunnel release surgery is performed to reduce the pressure in the region to arrest the progression of the disease. In previous studies, we have seen that carpal tunnel syndrome results in difficulty lifting and manipulating objects in day-to-day life. However, they are able to perform most tasks with some difficulty in an inefficient manner, due to the sensory deficits (Zhang et al., 2011, 2012).

Despite extensive work on CTS grasp, it remains unknown whether preparatory changes like ASAs are affected in CTS or recover after decompression. This constitutes a theoretical and clinical gap because synergy analysis can reveal neurological dysfunction even in the absence of overt symptoms (Lewis et al., 2016). Notably, anticipatory synergy adjustments (ASAs) have emerged as particularly sensitive indicators, detecting neurological impairments even when overall synergy indices appear unaltered (Jo et al., 2016). Taken together, these studies highlight the unique potential of ASA analysis to detect changes in motor coordination that may not be captured by conventional assessment approaches. Two parameters of ASA can be quantified: (i) ASA onset, defined as the earliest premovement time point at which the synergy index deviates below its baseline by a predefined threshold; and (ii) ASA amplitude, defined as the maximum drop in the synergy index from baseline before movement. Notably, these features are differentially impacted in various pathologies (Jo et al., 2016; Lewis et al., 2016).

Carpal tunnel release surgery selectively restores peripheral sensory conduction while leaving central motor circuits largely intact. This provides a unique opportunity to dissociate the contribution of sensory feedback to force regulation from the neural mechanisms underlying anticipatory coordination. While improved grip force control following decompression has been well documented, it remains unknown whether anticipatory synergy adjustments reflecting feedforward reorganisation of multi-digit coordination prior to a destabilising action are altered by changes in peripheral sensory input. If ASAs depend critically on sensory feedback, their timing or magnitude should change following surgery. Conversely, if ASAs primarily reflect supraspinal control processes, they should remain invariant despite improvements in force regulation. To test these competing predictions, we examined multi-digit coordination during a self-generated, destabilising manipulation task in individuals with carpal tunnel syndrome before and after surgical decompression, quantifying both force regulation and anticipatory synergy adjustments using variance-based analysis (Latash et al., 2002; Scholz & Schöner, 1999).

## 2. Methods

### Participants

Eleven participants (age: 54 ± 16 years; seven females and four males) were recruited between December 2022 and June 2024 for the study from the SRM Institute of Medical Sciences (SIMS) in Vadapalani and the Tamil Nadu Government Super Speciality Medical College (TNGSSMC) in Omandur, Chennai. All participants were right-handed; three had carpal tunnel syndrome (CTS) in their left hand, while the remaining eight had it in their right hand. The participants, who were scheduled for surgery, were selected based on clinical examinations and electrodiagnostic tests that confirmed the diagnosis of CTS. Exclusion criteria included individuals undergoing steroid therapy, those with significant finger rigidity, and participants with psychiatric or neurological impairments that might affect their ability to complete the experimental tasks. All participants provided written informed consent, and the study followed the guidelines of the Declaration of Helsinki. Ethical clearance was obtained from the institutional ethics committee at each of the participating study centres (SIMS: SIMS IEC/Others/41/2022; TNGMSSH: 1577/P&D-1/TNGMSSH/2023/BMS/O18).

### Experimental setup

The experimental setup consisted of an instrumented handle with force/torque sensors to measure the forces and moments of forces in 3 dimensions for each of the five fingers placed on the handle. A spirit level was attached to the top of the handle on the participant’s side for the participants to ensure that the handle is held correctly. On the opposite side of the top block, an inertial measurement unit (IMU: BNO055, Bosch, Germany) was attached to measure the orientation of the handle (Figure 1(A)). The IMU was configured to output quaternions for calculating the handle’s orientation. Prior to the beginning of a trial, the quaternions were recorded for one second and averaged to create a baseline reference quaternion (Q_ref_). During each trial, the quaternions measured (Q_trial_) were expressed relative to this reference using quaternion conjugate multiplication:

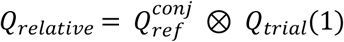

**Figure 1:**
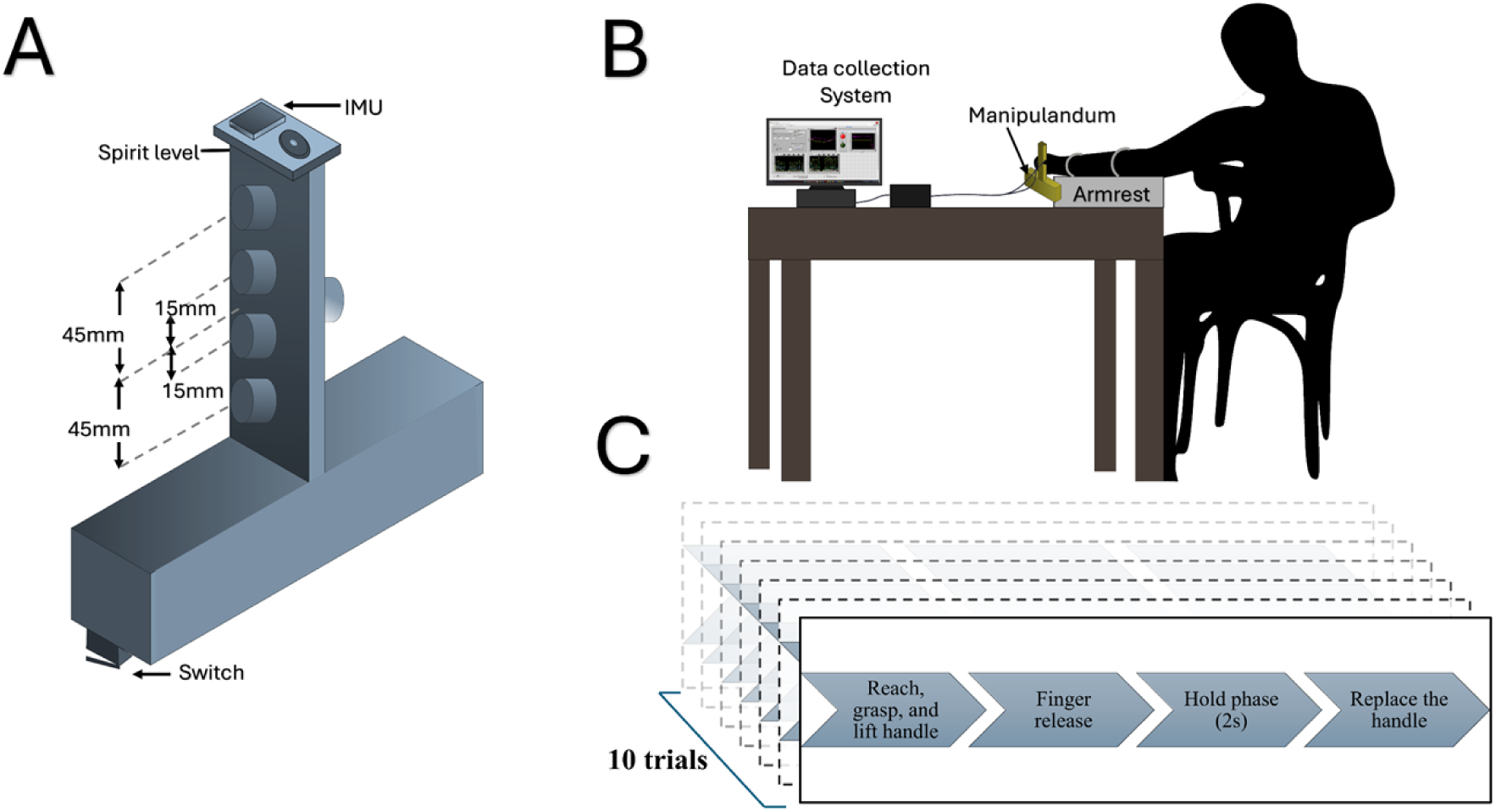
Experimental apparatus and protocol: **(A)** The figure shows the custom-designed instrumented handle used for multi-finger force and moment measurement during the grasp-and-lift task. Five force/torque sensors are attached corresponding to each finger contact location to capture three-dimensional fingertip forces and moments from the thumb, index, middle, ring, and little fingers. A spirit level atop the block (participant’s side) allowed subjects to maintain vertical alignment, while an inertial measurement unit (IMU) attached on the opposite side provided precise measurement of handle orientation in three-dimensional space via quaternion data. Two limit switches mounted at the base detected the exact time of handle lift-off. The total weight of the handle was approximately 800g. (B) Participants lifted the instrumented handle using all five fingers, maintaining vertical alignment. After holding it stable in the air, they were instructed to remove their index finger while keeping the handle steady with the remaining fingers, and to hold the object in this configuration for two seconds before replacing it on the table. (C)The timing diagram illustrates the sequence of events in the experimental trials.

These relative quaternions were then used to determine the overall tilt of the handle. Since the overall experiment depended on maintaining the handle in rotational equilibrium, any trial with a tilt exceeding 3° was excluded from analysis. Two limit switches were attached to the base of the handle to identify the lift onset, or the time when the handle is lifted off the table. The total mass of the handle was 800g. The force data was collected and synchronised with the rest of the setup using LabVIEW virtual instrumentation. The data was collected at 100Hz.

### Experimental paradigm

Participants were instructed to lift and hold the instrumented handle using all five fingers, maintaining vertical alignment as indicated by the spirit level. After achieving a stable hold as confirmed by the experimenter, they were asked to quickly remove their index finger while keeping the handle steady with the remaining fingers, and to hold the object in this configuration for two seconds before replacing it on the table. This rapid, trial-timed release is expected to provoke both anticipatory and compensatory coordination strategies.

A total of 10 trials were recorded. Baseline measurements (Pre-op) were obtained prior to surgery. Follow-up measurements (Post-op) were conducted at least 3 weeks after surgery, following wound healing and resolution of postoperative pain, and were scheduled to coincide with the first routine hospital visit (mean ± SD: 5 ± 1 weeks post-op; Figure 1C).

### Data processing

Each force-torque sensor generated a 6-channel data corresponding to the forces (F_x_, F_y_ and F_z_) in the 3D axes as well as the moments (M_x_, M_y_ and M_z_) about these axes for each of the five fingers. Additionally, the quaternion values (4 channels) were recorded from the IMU. Lift onset times were obtained by detecting the rising edge of the output signal from the limit switch. The time of finger release was captured as the time when the value of the index finger normal force fell to less than 10% of the maximum value. All post-processing on the collected data was done using MATLAB 2019b.

### Finger forces

A zero-lag, second-order Butterworth low-pass filter was applied to remove noise above 15 Hz. Grip forces were measured across three phases of the grasp: (1) the baseline phase, defined as the 500 ms static hold period before finger release; (2) the transient phase, defined as the 150 ms immediately following finger release; and (3) the steady-state phase, defined as the stabilised period after finger release. The steady-state phase measurement began 250 ms after finger release and was averaged over a 500 ms window.

### Compensatory changes in force

To quantify compensatory adjustments in the forces of the middle, ring, and little fingers in response to index finger release, the changes in their forces were measured relative to the change in index finger force. The change in index finger force was calculated as the difference between the mean force during the 500 ms before release and the mean force during the 500 ms after release. Similarly, the force changes in the other fingers were calculated and normalised with respect to the index finger force change. Consequently, the percentage change in finger forces relative to the index finger force drop was obtained for all trials across all participants. Percentage change data were used to normalise for inter-participant differences in baseline force magnitude, facilitating comparison across fingers and intervention.

### Moment compensation

The task required to maintain the object at rotational equilibrium during the finger release for maintaining the roll within the acceptable range. This could be calculated using the moment equation. Moment compensation was computed based on the equation:

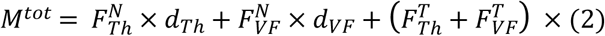

where, *M*^*tot*^ is the net moment applied on the handle, 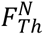 is the moment due to the thumb normal force, is the moment due to the net normal force from the virtual finger, or the mechanical equivalent of the four fingers (index, middle, ring and little) acting on the object; 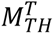 is the moment due to the tangential force at thumb sensor and 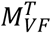 is the moment due to the net tangential force due to the virtual finger.

### Variance analysis for computing synergy

Variance analysis was done for the static hold phase of all trials for all participants to study multi-finger synergy (M. L. Latash et al., 2002). The synergy index (ΔV) was computed as the normalised difference in the variances of the task-dependent variable, referred to as the performance variable (PV) and the sum of variances of the individual elements (elemental variables or EV) contributing to the former, according to the following equation:

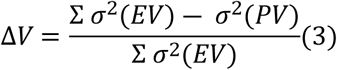

Given the multi-dimensional force–moment structure of the grasp-and-lift task, coordination was quantified using the synergy index (ΔV) without explicit decomposition into UCM and orthogonal variance components. In the current scenario, the synergy analysis is performed on the moment compensation task at the level of the thumb and the virtual finger.

The moment equation (1) can be rewritten as:

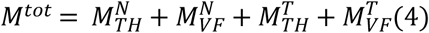

Where, the elements on the RHS of the equation are the elemental variables stabilising the net compensatory moment (performance variable).

The synergy index was computed across all trials of a participant. Strong negative covariation between the elemental variables indicates a synergy stabilising the performance variable and will result in a positive synergy index (ΔV). Poor synergies will result in values closer to zero, whereas purposeful destabilisation, sometimes referred to as anti-synergy, will result in negative synergy indices. Since the synergy index is a ratio, the index is transformed to a normal distribution using Fisher’s transformation for statistical analysis. This transforms the bounded distribution [-1,1] to a normal distribution (-∞,∞) according to the following formula:

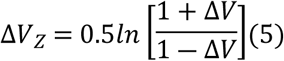

### Anticipatory synergy adjustments

Anticipatory synergy adjustments are defined as the change in the synergy index before a task in preparation for the task. We derive two metrics indicative of the synergy adjustments: (a) the onset of ASA and (b) the amplitude of ASA. The timing of ASA was calculated as 1 standard deviation dip in the synergy index from the baseline synergy index measured between 1000ms and 500ms before the finger release time (t_0_). To measure the amplitude of ASA, we took the mean synergy index during the baseline period (1000- 500ms before t_0_) and measured the minimum value of ASA with respect to it before finger release.

### Statistical analysis

A two-way repeated measures ANOVA was conducted to examine the effects of Intervention (Pre-op, Post-op) and Phase (Baseline, Transient, Steady State) on total grip force. Another two-way repeated measures ANOVA was conducted on percentage change in force data, focusing on the factors of Fingers (Middle, Little, and Ring) and Intervention (Pre-op and Post-op) to evaluate the variations in force between these conditions. Although bounded data can violate assumptions of ANOVA, distributions were checked for normality using the Shapiro-Wilk test, and ANOVA residuals did not deviate from normality, supporting use of parametric tests.

In addition, to investigate the changes in ASA due to surgery, a multivariate ANOVA was performed on the factor Intervention (2 levels: Pre-op and Post-op) for dependent variables, onset of ASA, as well as amplitude of ASA. Equivalence between two conditions was established by using the two one-sided t-tests (TOST) procedure for each variable with predefined standardised equivalence bounds of d_z_ = ±0.9 (SESOI, computed via GPower).

The significance level (α) was set at 0.05 for all statistical tests. The normality of the data was established using the Shapiro-Wilk test.

## 3. Results

### Task performance

The participants performed the task according to the provided instructions. They needed to keep the handle’s orientation aligned with the vertical axis within an acceptable range throughout the trial. Any trial in which the roll exceeded 3° during the finger removal phase was discarded. This step is crucial for ensuring meaningful results in the synergy analysis. The mean roll for pre-op trials was 1.21 ± 0.45° and was 1.55 ± 0.9° for post-op trials.

### Analysis of finger forces

The total grip force decreased after surgery for all three phases within the trial (Table 1). A two-way repeated measures ANOVA revealed a significant main effect of Intervention (F_(1,9)_ = 10.91, p <0.01, *η*^2^ =0.55), with participants exerting significantly lower grip forces following carpal tunnel release compared to pre-surgical trials (Table 1, Figure 2).

**Figure 2:**
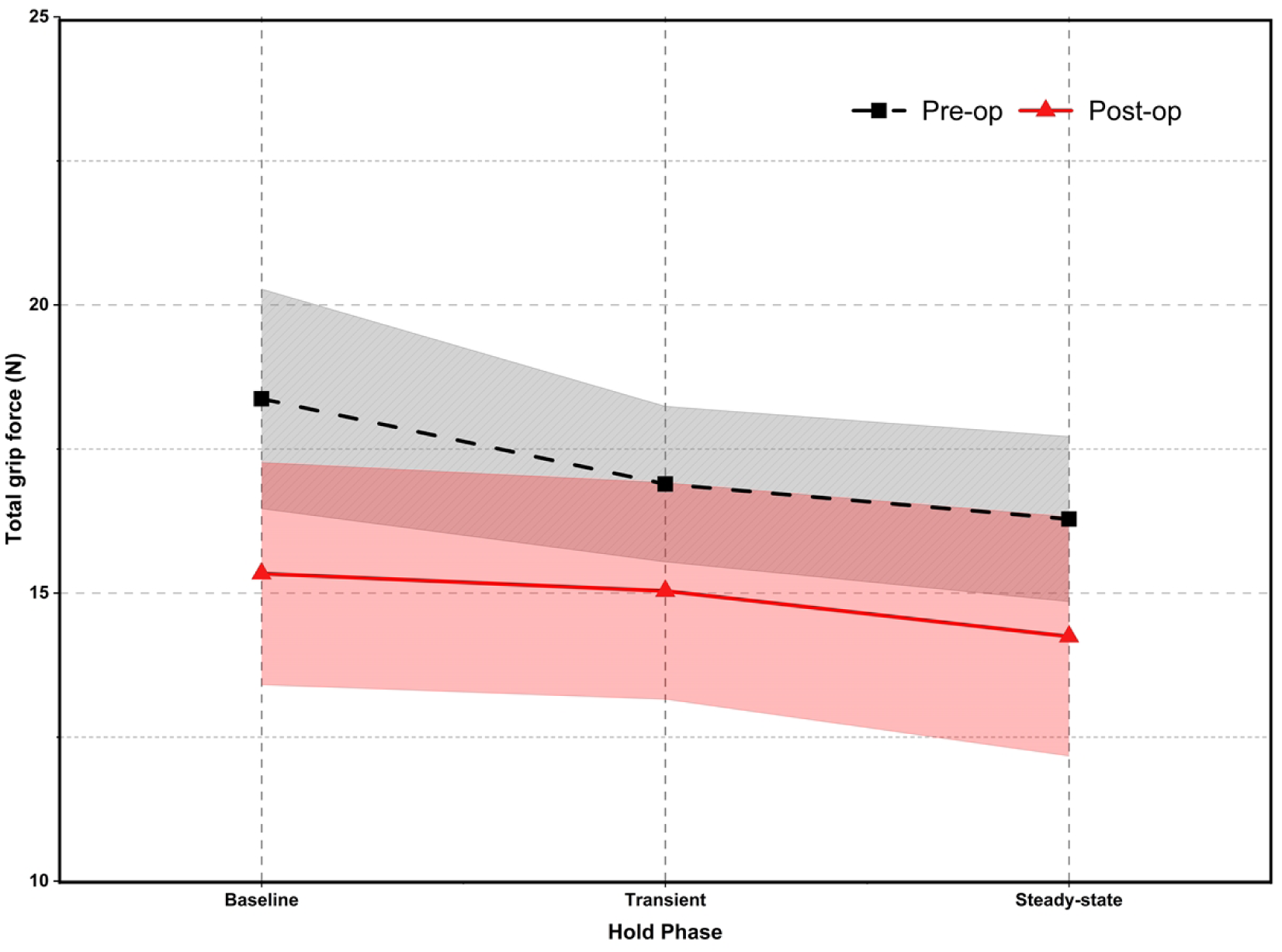
Reduction in total grip force across task phases before and after carpal tunnel release surgery: The line plot depicts changes in mean total grip force ( in Newtons) across three trial phases: baseline (static hold before finger release), transient (immediately after release), and steady-state (post-adaptation). The dashed black line represents preoperative (Pre-op) group means, while the solid red line denotes postoperative (Post-op) means. Shaded regions around each line show ±1 standard error. Each data point (square for Pre-op, triangle for Post-op) represents the phase mean for n = 11 participants. Grip force was consistently lower after surgery across all phases compared to preoperative values. This consistent reduction reflects more efficient grip force calibration and improved sensorimotor feedback following carpal tunnel decompression (see also Table 1 and Results for statistical analysis). The figure demonstrates that surgical intervention leads to overall improvement in grip force regulation during the experimental task.

**Table 1:** Total grip force (Mean ± SD) before and after surgery in each phase of the trial.

| Phase | Pre-op (Mean ±SD) | Post-op (Mean ±SD) |
| --- | --- | --- |
| Baseline | $18.37 \pm 1.90$ N | $15.34 \pm 1.93$ N |
| Transient | $16.89 \pm 1.35$ N | $15.04 \pm 1.88$ N |
| Steady-state | $16.28 \pm 1.43$ N | $14.25 \pm 2.07$ N |

Two-way repeated measures ANOVA was performed on percentage force change data for factors Fingers (Levels: Middle, Little and Ring) and Intervention (Levels: Pre-op and Post-op). There was no significant effect of intervention (pre-op mean: 19.34 ± 6.53% SEM; post-op mean: 28.54 ± 5.01% SEM) on the force sharing pattern. In contrast, there was a significant main effect of Finger on percentage force change (F_(1.3, 10.3)_ = 33.21, p <0.0001). The middle finger showed the highest mean change (69.25 ± 10.46%), followed by the ring finger (14.46 ± 4.99%), and the little finger (–11.88 ± 8.31%). Bonferroni-adjusted post hoc comparisons revealed significant differences between all pairs of fingers (p < 0.02) (Figure 3). The distribution of ANOVA residuals was assessed using the Shapiro-Wilk test and found to be approximately normal (p > 0.05), supporting the validity of the parametric test.

**Figure 3:**
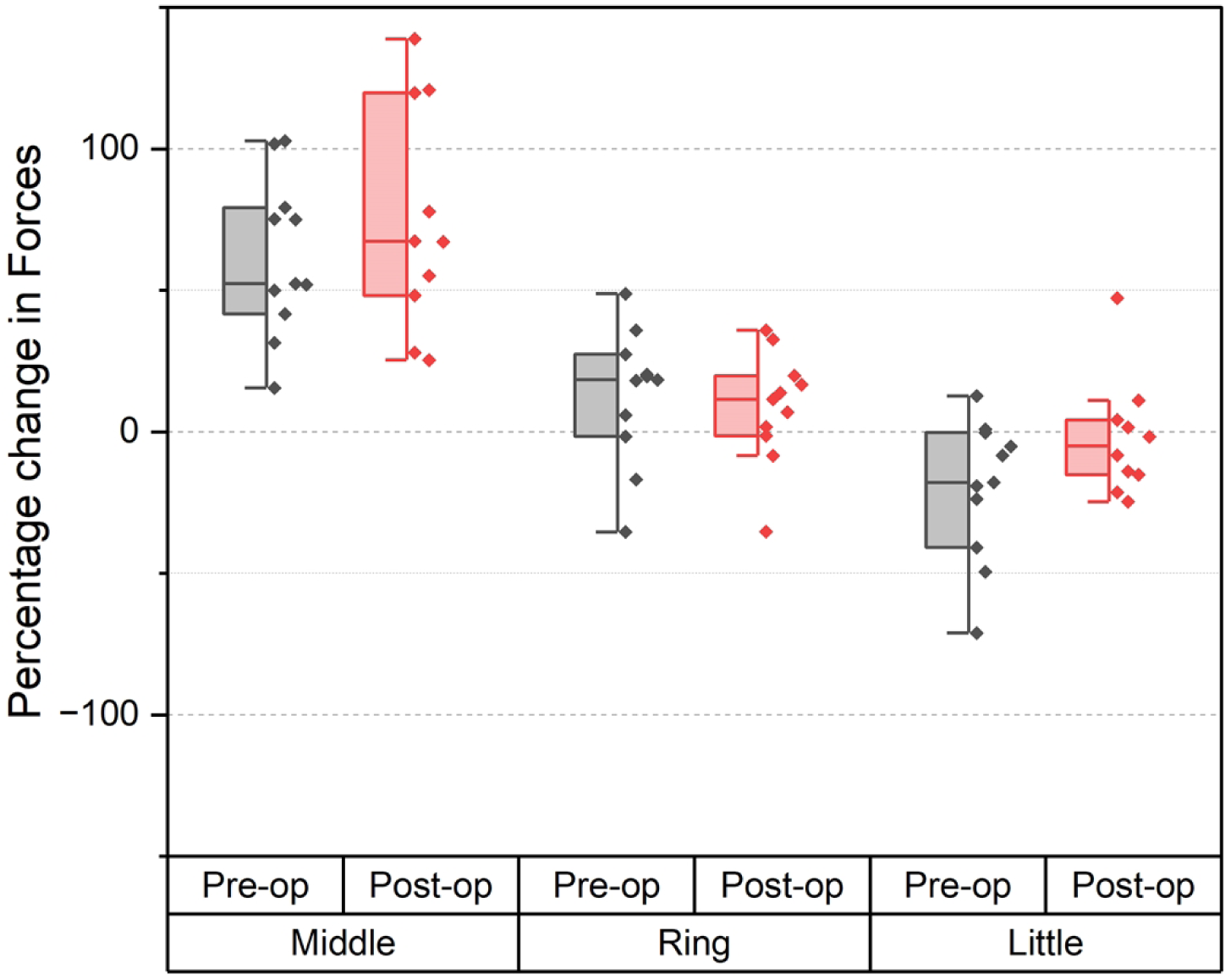
Percentage force compensation by middle, ring, and little fingers during index finger release pre- and post-surgery: Plots represent the finger force compensation as a percentage of the net drop in index finger force for each finger (middle, ring, little) before (pre-op) and after (post-op) intervention. Data are shown as mean ± standard error. The results are displayed using grouped half-box indexed plots, where each half box represents the distribution of finger force compensation values for a specific finger (middle, ring, little) within each intervention condition (pre-op and post-op). For each group, the box indicates the interquartile range, the line marks the median, and individual data points are shown alongside. The overall force-sharing strategy did not change with surgery (no effect of intervention, no intervention × finger interaction), whereas the middle finger consistently carried the largest compensatory load, followed by the ring, with the little finger contributing the least; all fingers differed from each other.

### Anticipatory synergy adjustments (ASA)

A multivariate ANOVA was performed for factor Intervention (Pre-op and Post-op) on dependent variables, Onset of ASA, as well as amplitude of ASA. No significant effect was observed for either variable after surgery. To test the similarity of groups, two one-sided t-test (TOST) analysis was done to establish that the two conditions were significantly similar with the smallest effect size of interest (SESOI =0.9) as the equivalence bounds.

The onset of ASA yielded nearly identical means before (-0.19±0.04) and after surgery (-0.17±0.04). TOST equivalence testing confirmed statistical equivalence, with t_(10)_ = 20.43, p < 0.001 for the lower bound and t_(10)_ = -19.54, p < 0.001 for the upper bound as shown in Figure 4. Likewise, the amplitude of ASA was equivalent pre-op (0.39±0.1) and post-op (0.58±0.1). This was confirmed by the TOST equivalence test ( t_(10)_ = 7.67, p < .001 (lower bound), t_(10)_ = -4.96, p < .001 (upper bound)) within equivalence margin, as shown in Figure 5.

**Figure 4:**
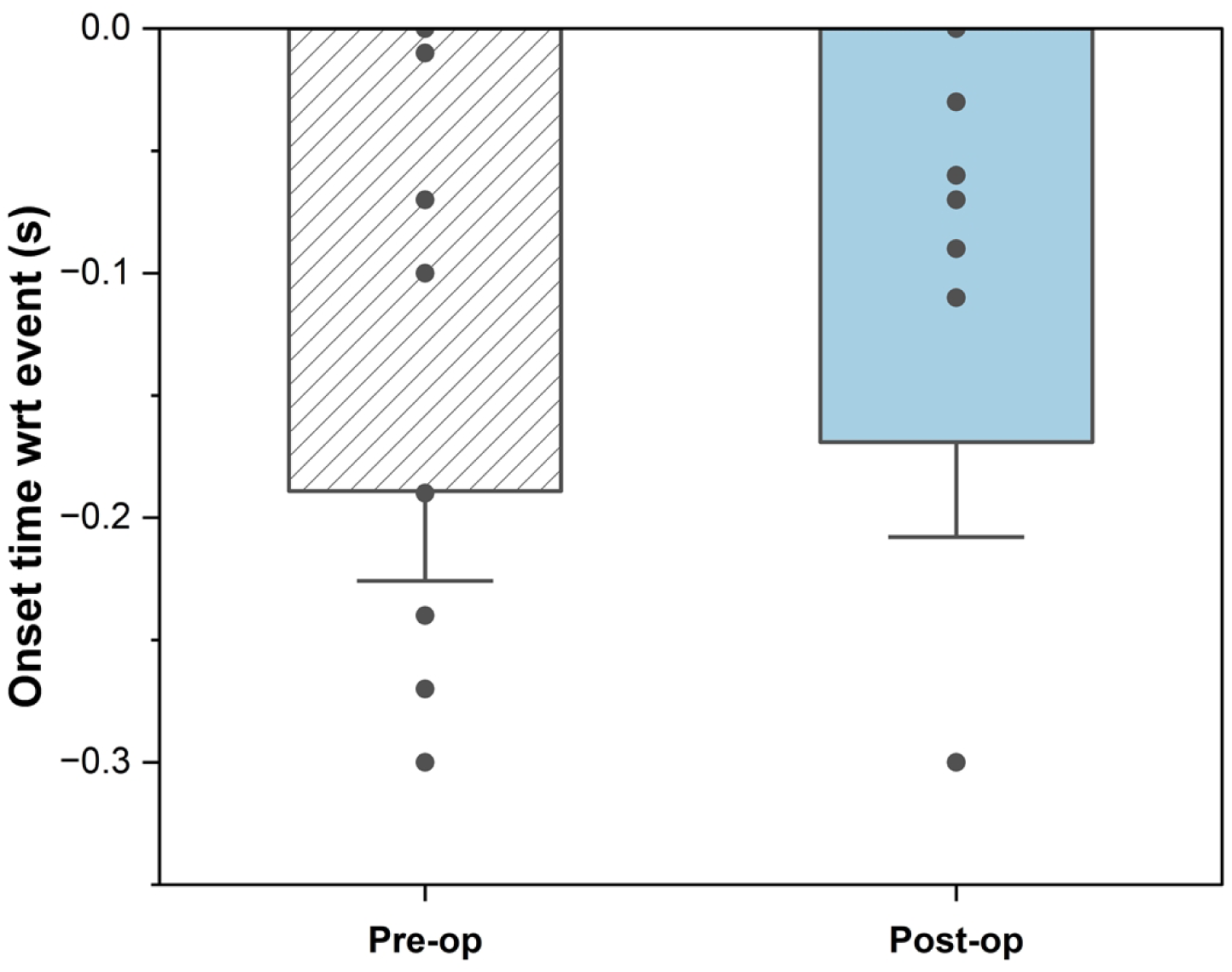
Onset timing of anticipatory synergy adjustments (ASA): Onset timing of ASA relative to finger release, shown pre- and post-surgery. Boxes indicate group distributions with individual participant data overlaid (n = 11). ASA onset timing did not show a systematic change following carpal tunnel release.

**Figure 5:**
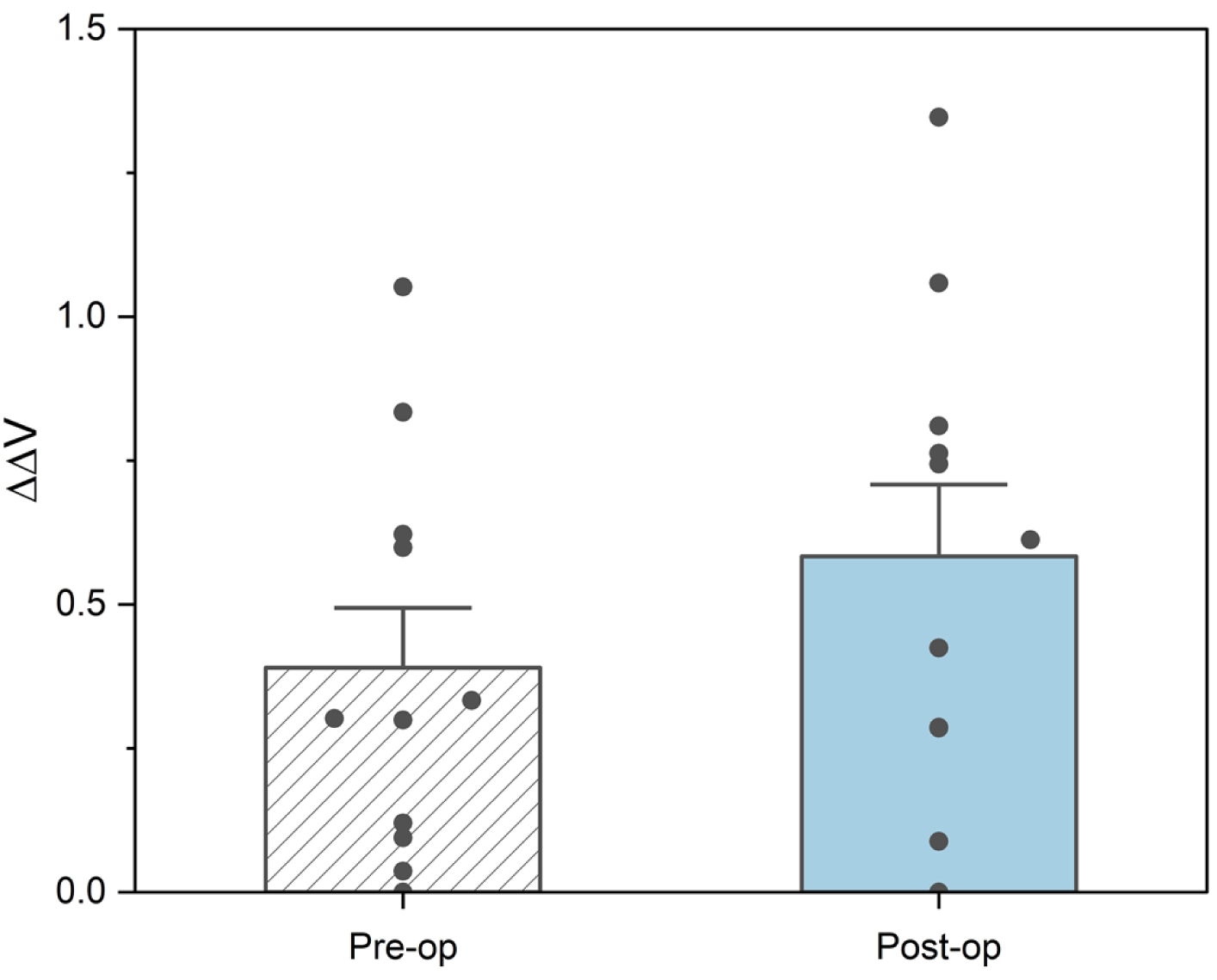
Amplitude of anticipatory synergy adjustments (ASA): Amplitude of ASA during the static hold period before movement onset, shown pre- and post-surgery. Boxes indicate group distributions with individual participant data overlaid (n = 11). ASA amplitude did not show a systematic change following carpal tunnel release.

## 4. Discussion

This study examined the effects of carpal tunnel release (CTR) surgery on fingertip force coordination, focusing on compensatory mechanisms and anticipatory synergy adjustments during self-induced perturbations in a grasping task. We expected an overall improvement measured in terms of fingertip forces, compensatory mechanisms, and anticipatory synergy adjustments. Fingertip forces improved across all trial phases following surgery, as reflected by lower and more optimal force levels compared to preoperative values. However, compensatory mechanisms, measured by changes in the forces exerted by intact fingers to offset the removal of a digit, remained similar before and after surgery. Additionally, the timing and amplitude of anticipatory synergy adjustments (ASAs) before and after surgery were statistically equivalent, despite improvements in sensory function and static force control. The following paragraphs discuss these results in detail.

### Reduced Grip Force: Enhanced Sensorimotor Feedback

A significant reduction in total grip force was observed across all task phases following carpal tunnel release (see Table 1 & Figure 2), indicating that the surgical intervention was effective. This reflects improved sensory feedback, allowing for more precise force calibration, as hypothesised. Prior to surgery, impaired sensation due to median nerve compression typically leads to higher grip forces as a compensatory strategy to prevent slipping and maintain a stable hold (Zhang et al., 2011). Experimental median nerve compression in healthy participants also leads to increased grip force, as documented by Cole et al. (2003), further emphasising the role of tactile feedback in grip modulation. As sensory conduction improves, tactile feedback is efficiently utilised for better force control, as shown in clinical studies on postoperative changes in CTS (Hsu et al., 2009; Nishimura et al., 2003).

Unlike the stability observed in the initial hold and final steady state, the transient phase presents an unstable scenario characterised by the sudden removal of a finger. Despite the mechanical challenge introduced by finger removal, the transient phase still showed well-controlled force distribution with rapid adaptation by the remaining fingers. In the transient phase, force adaptation was controlled rather than maintaining pre-removal force levels.

### Compensation Patterns in Finger Force Redistribution

When the index finger was removed, most of the compensatory load was taken by the middle finger, followed by the ring and little finger, respectively. This effect of proximity aligns with previous studies indicating a diminishing compensatory role with increasing distance from the perturbed digit (Jacob et al., 2022). The compensatory force-sharing pattern among the remaining fingers in response to the disengagement of the index finger from the grasped handle did not change after the surgery. The stability of multi-finger force distribution observed after surgery may reflect the persistence of robust underlying neuromechanical and neural linkages, which support coordination even after peripheral alteration.

Two primary mechanisms govern inter-finger interactions: enslaving and error compensation. Enslaving describes the involuntary co-activation of neighbouring fingers, driven by shared neural pathways and biomechanical linkages, which produces positively correlated forces between the perturbed digit and its neighbours (Kilbreath S L & Gandevia S C, 1994; Zatsiorsky et al., 2000). In contrast, error compensation reflects the active, negative correlation of forces among fingers as the nervous system modulates recruitment to maintain grip stability when the contribution of some of the fingers changes (Kruger et al., 2007). The interplay between these mechanisms could be understood from two different studies that employed distinct perturbation paradigms. In the inverse saxophone study, which required participants to grasp and lift an object with a changing finger-thumb distance, enslaving effects dominated the response of unperturbed fingers, particularly in proximal digits (Jacob et al., 2022). In contrast, the inverse piano paradigm, where participants maintained constant total force while one finger was displaced, revealed compensatory reductions in neighbouring finger forces, emphasising error compensation (Martin et al., 2011). Likewise, our current task, necessitating stability during finger release from the grasped handle, displays compensatory adjustments that dominate over enslaving effects, suggesting that task goals critically shape inter-finger responses.

Santello et al. (2013) provided a synergy framework for such control, describing two temporal components coordinating multi-digit actions. The common drive synergy produces in-phase force changes across fingers to ensure gross grip stability, whereas the error compensation synergy facilitates out-of-phase, reciprocal modulations for finer adjustments. These dual mechanisms offer a conceptual bridge between observed coordination patterns: the former may reflect enslaving effects, while the latter is responsible for behaviour that overrides enslavement in favour of error compensation.

This interpretation is further corroborated in electromyographic studies. Maier & Hepp-Reymond (1995) reported minimal motor unit synchronisation during precision grip tasks, consistent with individuated finger control and the influence of error compensation. In contrast, Huesler et al. (1998) observed increased synchronisation during power grips, indicative of more unified muscle activation patterns, which align with the common drive synergy described by Santello. Together, these findings underscore that the neural balance between enslaving and compensatory mechanisms is not fixed, but rather shaped by task demands and context.

Building on this, Martin et al. (2008) established that error compensation can dominate and mask enslaving effects at higher force levels, demonstrating a task-dependent shift in coordination strategies. In our study, the unchanged compensatory patterns post-surgery likely reflect this stable integration of mechanisms, modulated by task demands rather than peripheral structural changes. This suggests that neuromechanical coupling in the hand is robust and subtly balanced, maintaining force-sharing synergies despite anatomical perturbations.

### Anticipatory surgery adjustments unaffected by surgery

This study confirmed that anticipatory synergy adjustments (ASA), specifically onset timing and amplitude of ASA, remained unchanged in patients with carpal tunnel syndrome (CTS) after carpal tunnel release (CTR) surgery. Previous studies have consistently demonstrated that ASAs represent a crucial feed-forward control mechanism that prepares the motor system for upcoming perturbations by transiently reducing synergy strength 100-300 ms before movement onset (Jae et al., 2005; Olafsdottir et al., 2005). The preservation of ASA timing and amplitude in our CTS patients suggests that this anticipatory mechanism may be more resilient to peripheral sensory deficits than previously thought.

This finding contrasts sharply with other studies on different populations. Studies of Parkinson’s disease (PD) patients have consistently shown both delayed ASA onset and reduced amplitude compared to healthy controls (Jo et al., 2015; Park et al., 2012). Similarly, patients with multiple sclerosis (MS) demonstrate significantly delayed ASA initiation and smaller magnitude changes (Jo et al., 2017). Even healthy elderly individuals show delayed and reduced ASAs compared to younger adults (Olafsdottir et al., 2007, 2008). This is mechanistically plausible, as both PD and MS produce direct disruptions in central neural mechanisms critical for anticipatory feedforward planning and motor coordination. In PD, degeneration of basal ganglia circuitry impairs the initiation and temporal precision of feedforward adjustments (Jiang, 2022), whereas in MS, demyelination and white matter lesions similarly disrupt central motor pathways (Lubetzki & Stankoff, 2014). Ageing, in turn, is associated with global neurodegeneration that affects both central and peripheral pathways, further slowing anticipatory coordination and reducing its magnitude (Nieto-Sampedro & Nieto-Díaz, 2005).

One possible explanation for the absence of ASA modulation in the present study is that changes in anticipatory coordination may emerge only over longer recovery intervals. However, evidence from PD indicates that both deterioration and improvement of ASA parameters can occur over relatively short intervention periods, such as during withdrawal and reinstatement of dopaminergic therapy (Park et al., 2012). If peripheral sensory deficits in CTS exerted a strong influence on feedforward control, at least modest changes in ASA timing or magnitude would be expected within 3-5 weeks following decompression. The absence of such changes in the present data, therefore, suggests that anticipatory feedforward coordination remains largely preserved in CTS during early recovery.

This interpretation is further supported by recent work using a multi-finger force production task, which reported preserved anticipatory synergy adjustments in individuals with CTS despite weakened force-stabilising synergies (Abolins et al., 2026). Together, these findings point to a dissociation between feedback-dependent force regulation and feedforward anticipatory coordination, with the latter showing relative robustness to peripheral sensory disruption and restoration.

These findings have important theoretical implications for our understanding of the neural basis of anticipatory control. The preservation of ASAs in CTS patients supports the hypothesis that these adjustments rely primarily on central feed-forward mechanisms rather than peripheral sensory feedback loops (Houk, 2005). This is consistent with the central back-coupling hypothesis, which proposes that synergy adjustments are mediated by internal neural networks that can function independently of immediate sensory input (Goodman & Latash, 2006; Latash et al., 2005; Naik et al., 2024). In this framework, CTS primarily alters peripheral sensory input, whereas PD, MS, and ageing involve central neural degradation that directly compromises predictive motor programs. Consequently, peripheral sensory restoration alone may be insufficient to modify anticipatory coordination.

The present findings also extend our earlier work demonstrating preserved synergy indices during static grip following carpal tunnel release (Jacob et al., 2025). While static synergy indices and ASAs are related, prior studies indicate that they can be differentially affected by neurological pathology. For example, patients with subcortical disorders often show both reduced synergy indices and diminished ASAs, whereas individuals with mild cortical stroke may exhibit preserved synergy indices alongside impaired ASAs (Jo et al., 2016; Lewis et al., 2016). The current study extends this dissociation to dynamic transitions during grasping, demonstrating that anticipatory planning of multi-digit coordination remains stable even under changing biomechanical constraints.

The study did not include a matched healthy control group, as the primary objective was to examine within-subject changes following surgical decompression, with each participant serving as their own baseline. Consequently, the findings address modulation of anticipatory coordination across recovery rather than deviation from normative performance. Assessments were conducted during the early postoperative phase (average ∼5 weeks), when participants had resumed hand use; therefore, longer-term adaptations in coordination cannot be excluded. In addition, a single dynamic transition (five-to-four-finger grasp) was employed to isolate anticipatory synergy structure under controlled conditions. While this design minimized task-related confounds, future studies should evaluate additional grasp transitions and extended recovery intervals to determine the generality and temporal evolution of these effects.

## 5. Conclusion

This study probed the effect of carpal tunnel release surgery on anticipatory control in preparation for a known task. Our findings emphasise that the brain’s core strategies for preparing and coordinating hand movements are robust to peripheral changes. The central nervous system maintains these anticipatory control mechanisms, despite changes in peripheral sensory input. Feedback from improved sensation after surgery served mainly to fine-tune grip efficiency rather than restructuring anticipatory control. Thus, functional recovery in CTS appears to be driven more by improved force regulation than by changes in predictive motor programs.

## Acknowledgement

We thank all the participants for their cooperation in this study. We are grateful to our colleagues at the BRAIN Lab at IIT Madras for their assistance with data collection. Additionally, we appreciate the support from the collaborating institutions for providing resources and facilitating our research.

